# RIP1 kinase activity promotes steatohepatitis through mediating cell death and inflammation in macrophages

**DOI:** 10.1101/2020.01.07.895516

**Authors:** Liang Tao, Yuguo Yi, Yuxin Chen, Haibing Zhang, Jiapeng Jie, Weigao Zhang, Qian Xu, Yang Li, Pontus Orning, Egil Lien, Mengshu Zhao, Pingshi Gao, Ling Ling, Zhao Ding, Chao Wu, Qiurong Ding, Junsong Wang, Jianfa Zhang, Dan Weng

## Abstract

Hepatocyte cell death and liver inflammation have been well recognized as central characteristics of nonalcoholic steatohepatitis (NASH), however, the underlying molecular basis remains elusive. The kinase receptor-interacting protein 1 (RIP1) is a key regulator of apoptosis, necroptosis and inflammation, we thus hypothesized that the kinase activity of RIP1 may be involved in the pathogenesis of NASH. Wild-type and RIP1 kinase-dead (*Rip1^K45A/K45A^*) mice were fed with methionine-and choline-deficient diet (MCD) or high-fat diet (HFD) to establish distinct NASH models. In both models, compared to WT mice, *Rip1^K45A/K45A^* mice exhibited significantly less liver injury, less steatosis, decreased inflammation, and less cell death in liver tissue. Moreover, hepatic fibrosis as characterized by Sirius Red staining, expression of α-SMA and other fibrosis markers, were significantly alleviated in *Rip1^K45A/K45A^* mice than WT controls. Furthermore, using bone marrow transplantation to create chimeric mice, we found that it is the RIP1 kinase in hematopoietic-derived macrophages contributing mostly to the disease progression in NASH. Results from *in vitro* studies were in agreement with the *in vivo* data, demonstrating that RIP1 kinase was required for inflammasome activation and cell death induced by saturated fatty acid (palmitic acid) in bone marrow-derived macrophages (BMDMs). At last, we also found that the phosphorylation and expression of RIP1 was obviously increased in patients with NAFLD or NASH, but not in healthy controls. In summary, our results indicate that RIP1 kinase is activated during the pathogenesis of steatohepatitis, and consequently induces inflammation and cell death in macrophages, contributing to the disease progression. Our study suggests that macrophage RIP1 kinase represents a specific and potential target for the treatment of NASH.

## Introduction

In the recent decades, the incidence of nonalcoholic fatty liver disease (NAFLD) increases rapidly and becomes a huge threat to public health of the whole world. Nonalcoholic steatohepatitis (NASH) is the aggressive subtype of NAFLD and is becoming the main cause for liver cirrhosis, hepatocellular carcinoma or liver transplantation in USA (1–3). Regarding the mechanisms involved in the pathogenesis of NASH, it is now widely recognized that hepatic lipid accumulation and increased free fatty acids (FFA) causes lipotoxicity to hepatocytes and induces cell death, which is accompanied by the massive recruitment of monocytes into liver and the occurrence of chronic inflammation (4–6). Accumulating studies suggest that hepatic cell death and inflammation play critical roles in the transformation of simple steatosis to steatohepatitis, promoting liver fibrosis and disease progression (7, 8). However, the molecular basis regulating hepatic cell death and inflammation in steatohepatitis still remains unclear.

The serine/threonine kinase RIP1, as a key regulator determining the cell fate, is a multitasking molecule harboring distinct roles with its kinase activity and kinase-independent scaffolding function(9, 10). In contrast to mediating NF-κB activation and cell survival through its scaffolding function, the kinase activity of RIP1 is critical in regulating apoptotic and necrotic cell death as well as inflammation (11). Activation of RIP1 kinase promotes the occurrence of either apoptosis or necroptosis, depending on the cellular context and the molecules it interacts. If activated RIP1 interacts with caspase-8, it leads to apoptotic cell death. When caspase-8 is deficient or inhibited, RIP1 interacts with RIP3 and MLKL to initiate necroptosis (12, 13). In addition to its key roles in mediating cell death, RIP1 kinase has also been found to mediate inflammatory pathways including inflammasome activation induced by different stimulus (14–16). Due to its important roles in regulating apoptosis, necroptosis and inflammation, which are all critically involved in the pathogenesis of NASH, we speculated that the kinase activity of RIP1 might also contribute to the pathology progression of NASH.

In this study, we comprehensively investigated the role of RIP1 kinase in the pathogenesis of NASH and the underlying mechanisms by comparing RIP1 kinase-dead knock-in mice and wild type mice in two different murine models of NASH. More importantly, using irradiation and bone marrow transplantation, our results also revealed that it is the RIP1 kinase activity in hematopoietic-derived macrophages contributing to the inflammation and fibrosis progression in NASH. In addition, using clinical samples, we indicated that the expression and activation of RIP1 was positively correlated with the pathology in NASH patients. Our results collectively demonstrated the functional involvement of macrophage RIP1 kinase activity in NASH development.

## Results

### RIP1 kinase activity prompted the severity of MCD-induced steatohepatitis

To investigate the role of RIP1 kinase in the development of steatohepatitis, RIP1 kinase-dead knock-in (*Rip1^K45A/K45A^*) mice, which contain a lysine point mutation (K45A) in the catalytic triad residues of the kinase domain as previously described (17, 18), were utilized. Wild type C57BL/6 and *Rip1^K45A/K45A^* mice were fed with normal chow as control (normal diet, ND) or MCD diet for 5 weeks to stimulate steatohepatitis. First, we examined whether RIP1 kinase was activated during MCD-induced steatohepatitis. As illustrated in Fig. 1A, the phosphorylation of RIP1 was obviously increased in the liver of MCD-fed wild type mice, but not in RIP1 kinase-dead mice. Next, we examined the liver damage, steatosis as well as fibrosis in different groups of mice. Compared to WT control, *Rip1^K45A/K45A^* mice exhibited significantly lower serum levels of alanine aminotransferase (ALT) (p<0.01) and hepatic triglycerides concentrations (p<0.05) (Fig. 1B&C). Histological analysis by hematoxylin and eosin (H&E) staining and Oil Red O staining demonstrated that the hepatic steatosis and lipid accumulation were markedly alleviated in *Rip1^K45A/K45A^* mice (Fig. 1D&E). MCD feeding significantly increased the transcriptional expression of α-smooth muscle actin (*α-sma*), which was much higher in WT mice than *Rip1^K45A/K45A^* mice (p<0.01) (Fig. 1F), suggesting that hepatic fibrosis was also attenuated in RIP1 kinase-dead mice. Taken together, these results suggest that RIP1 kinase was activated and the kinase activity of RIP1 was required to prompt the pathological severity of MCD-induced steatohepatitis.

**Fig. 1.**
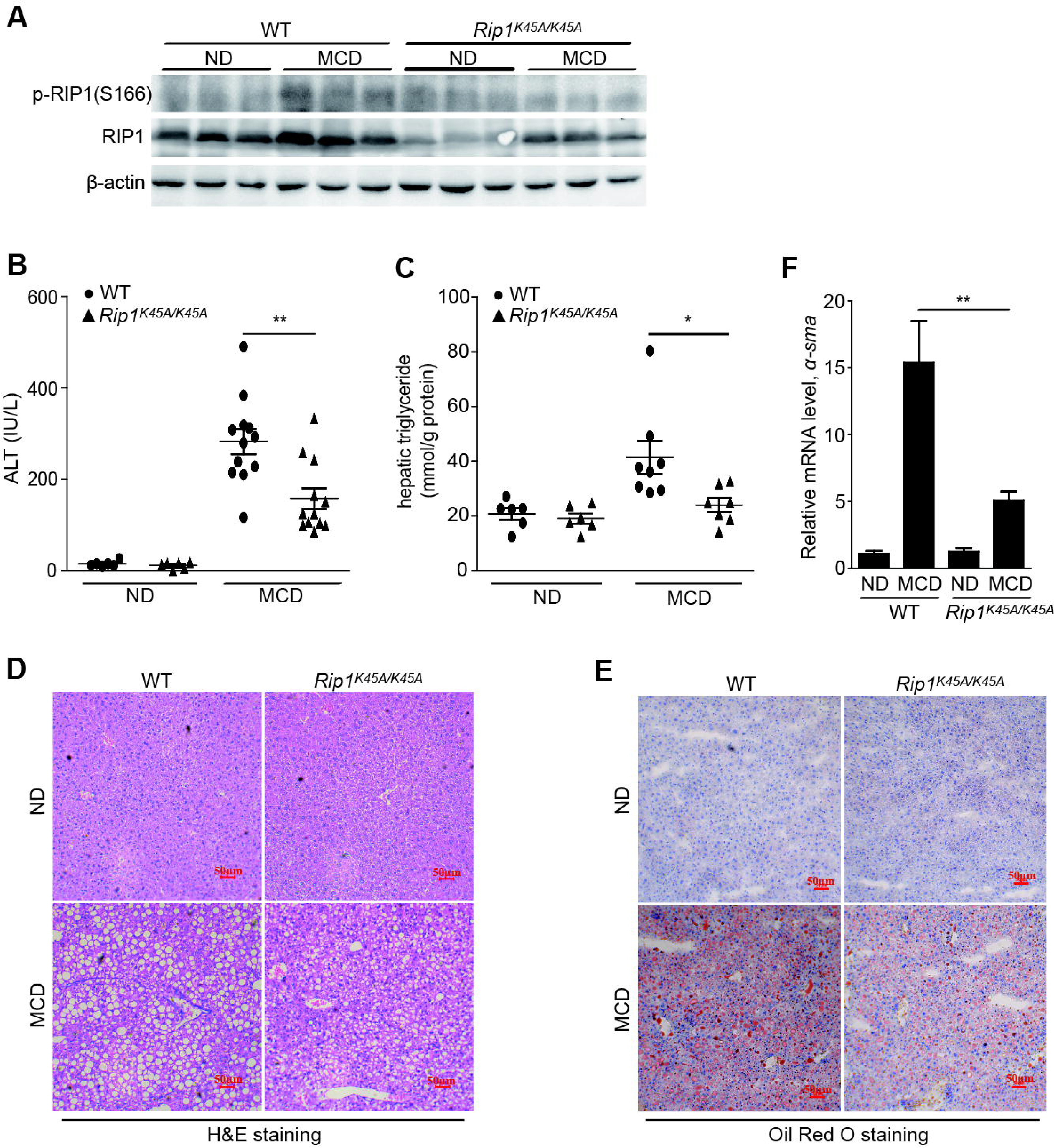
RIP1 kinase aggravated the severity of MCD-induced steatohepatitis. (A) MCD feeding induced the activation of RIP1 kinase as indicated by p-RIP1(S166) western blot in liver tissue of WT mice. (B) Serum ALT and (C) hepatic TG levels in WT and *Rip1^K45A/K45A^* mice treated with a normal diet (ND) or MCD diet. Representative images of (D) H&E staining and (E) Oil Red O staining of liver sections. (F) *α-sma* mRNA expression in the liver tissues of WT and *Rip1^K45A/K45A^* mice. Data are expressed as mean ± SEM (n=6 or 12 per group). ns not significant, *\*p*<0.05, *\*\*p*<0.01, \*\*\**p*<0.001.

### RIP1 kinase activity prompted the liver injury, steatosis and fibrosis in HFD-induced steatohepatitis

To further verify the role of RIP1 kinase activity in the development of NASH, another mechanistically distinct murine model of steatohepatitis induced by high-fat diet (HFD) feeding was used. Wild type C57BL/6 and *Rip1^K45A/K45A^* mice were fed with control diet or high-fat diet for 24 weeks. Body weight, food and water consumption, fasting blood glucose, glucose tolerance test (GTT) and basic metabolism parameters were monitored during HFD feeding. There was no obvious difference regarding the food/water intake, oxygen consumption, CO2 production and energy expenditure features between WT and *Rip1^K45A/K45A^* mice (Fig. S2), suggesting that RIP1 kinase inactivation did not affect the basic metabolism activity. For the body weight, although the initial body weight gain was lower in RIP1 KD mice than wild type controls, there was no significant difference for the final body weight at the end of HFD feeding between WT and RIP1 KD mice (Fig. S2A). Moreover, fasting blood glucose levels as well as glucose tolerance were similar between WT and *Rip1^K45A/K45A^* mice (Fig. S2D), suggesting that RIP1 kinase inactivation did not affect the development of obesity and insulin resistance in this dietary model. In contrast, both the serum ALT and hepatic TG levels were significantly reduced in *Rip1^K45A/K45A^* mice than WT following 24 weeks of HFD feeding (Fig. 2A-B). This was consistent with the MCD-NASH model. Reduction of steatosis in *Rip1^K45A/K45A^*mice was further confirmed by H&E and Oil Red O staining (Fig. 2C&D). Moreover, as indicated by Sirius Red staining, α-SMA immunohistochemistry analysis as well as the mRNA levels of transforming growth factor-beta1 (Tgf-β1), Col3a1 and α-sma, all results consistently demonstrated that HFD feeding induced obvious hepatic fibrosis in WT mice, but not in *Rip1^K45A/K45A^*mice (Fig. 2E-G). These results suggested that the kinase activity of RIP1 contributes to pathology development especially fibrosis progression in HFD-induced steatohepatitis, with no affect on food intake and obesity induction.

**Fig. 2.**
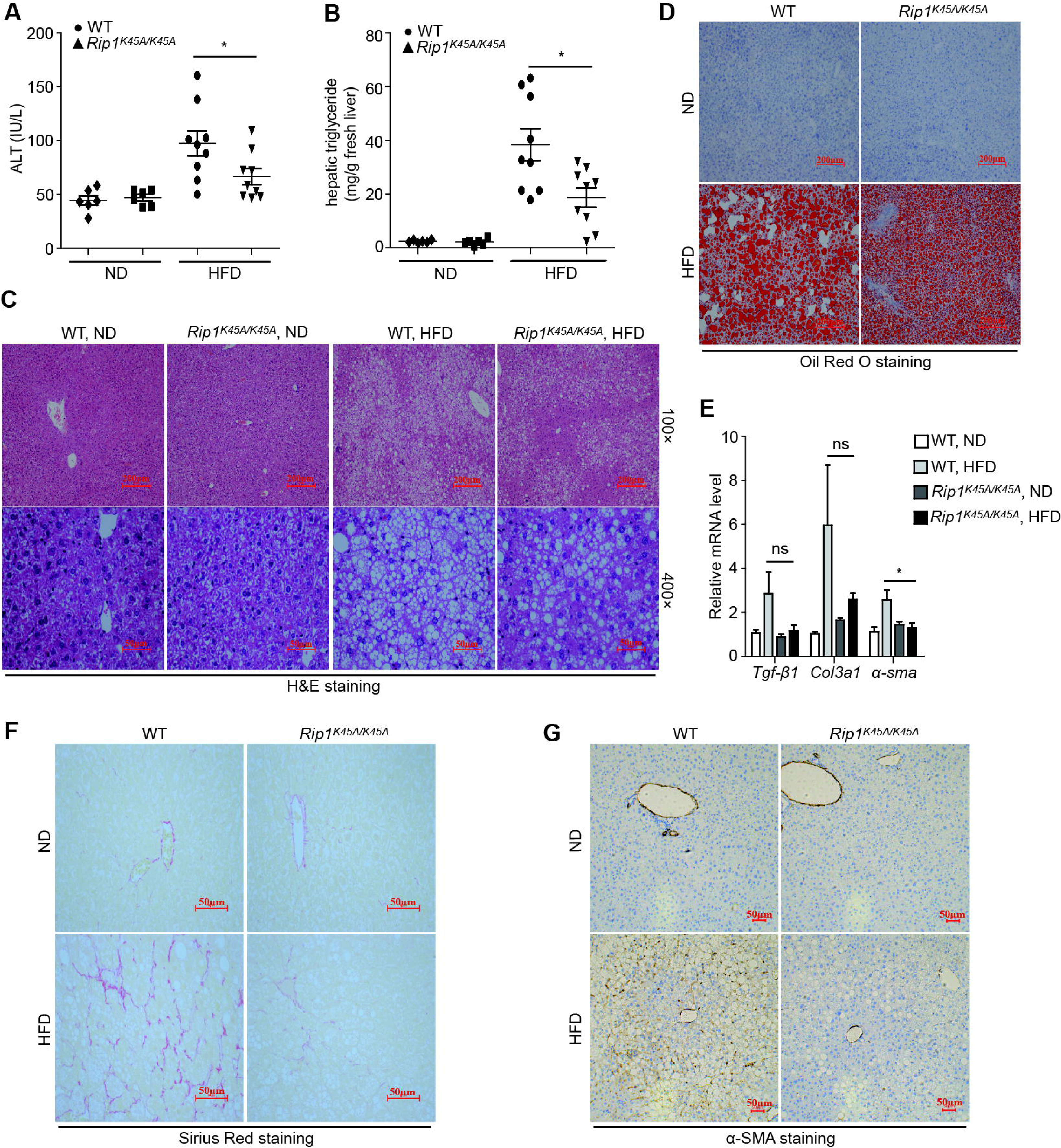
RIP1 kinase prompted the liver injury, steatosis and fibrosis in HFD-induced steatohepatitis. WT and *Rip1^K45A/K45A^* mice were fed with a control normal diet (ND) or a high-fat diet (HFD) for 24 weeks. Then (A) serum ALT and (B) hepatic TG levels were measured. Liver sections were analyzed with (C) H&E staining or (D) Oil Red O staining. Representative images of each group are presented. (E) Hepatic mRNA levels of *Tgf-β1*, *Col3a1* and *α-sma* in indicated groups. Representative images of (F) Sirius Red staining and (G) α-SMA protein expression as detected by immunohistochemistry in indicated groups. Data are expressed as mean ± SEM (n=6 or 9 per group). ns not significant, *\*p*<0.05, *\*\*p*<0.01, \*\*\**p*<0.001.

### RIP1 kinase activity contributed to the inflammation and hepatic cell death in both models of steatohepatitis

The above results suggest that the kinase activity of RIP1 is required for NASH pathogenesis in both MCD- and HFD-induced models. To explore the underlying mechanisms, since liver cell death and inflammation are concomitant key features of many liver diseases including steathohepatitis and RIP1 kinase has been identified as a critical regulator of cell death as well as inflammation, we therefore examined these features in liver tissues of both models. As shown in Fig. 3A and Fig. 4A, both MCD and HFD feeding induced obvious cell death in WT liver tissue, and the cell death rate was significantly reduced in *Rip1^K45A/K45A^*mice (Fig. 3B and Fig. 4B), as determined by TUNEL staining of liver sections. F4/80 immunohistochemistry analysis indicated that more macrophages were recruited into liver in WT mice than *Rip1^K45A/K45A^*mice during both MCD and HFD-induced steatohepatitis (Fig. 3D and Fig. 4D). Consistently, the mRNA levels of Mcp-1, TNFα, IL-6 and IL-1β, all of which are typical inflammatory markers, were markedly increased in the liver of MCD-fed WT mice than control diet-fed mice, and the increase was significantly inhibited in *Rip1^K45A/K45A^* mice (Fig. 3C). We also detected the activation of caspase-1 and IL-1β in liver tissues using western blot. Pro-caspase-1 is cleaved and activated by inflammasome complexes, and active caspase-1 will further lead to the maturation of pro-inflammatory cytokines IL-1β and IL-18 (19–21). Both cleaved caspase-1 and mature IL-1β were reduced in *Rip1^K45A/K45A^* liver compared to WT controls following MCD diet administration (Fig. 3E). This was consistent in HFD model, as both the transcriptional expression of IL-1β, TNFα, Mcp-1 (Fig. 4C) and the protein expression of IL-1β in liver tissues (Fig. 4E) were obviously reduced in *Rip1^K45A/K45A^* mice than WT control. These results demonstrated that both cell death induction and inflammation were significantly inhibited in *Rip1^K45A/K45A^* mice, and it correlated well with the attenuated pathology of steatohepatitis in different NASH animal models, suggesting that the kinase activity of RIP1 contributes to the pathogenesis of NASH via regulating cell death and inflammation.

**Fig. 3.**
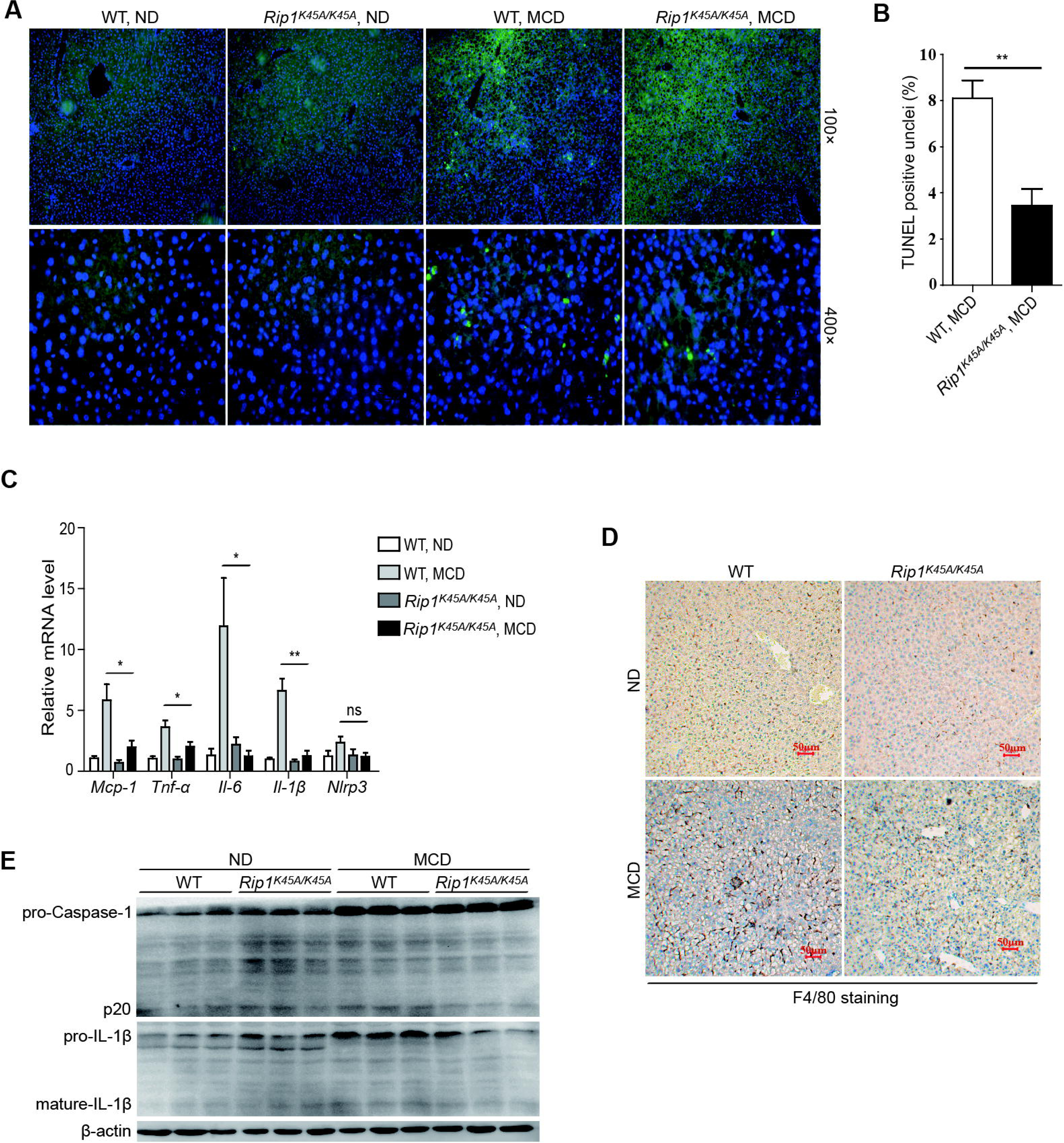
RIP kinase prompted the cell death and inflammation in the liver tissue of MCD-fed mice. (A) Representative TUNEL-stained liver sections and (B) quantification of the corresponding TUNEL-positive cells of WT and *Rip1^K45A/K45A^* mice. (C) Hepatic mRNA levels of the inflammatory molecules *Mcp-1*, *Tnf-α, IL-6, IL-1β* and *Nlrp3* in indicated groups. (D) Representative images of F4/80 analysis by immunohistochemistry staining in WT and *Rip1^K45A/K45A^* mice. (E) Immunoblotting detection of the pro-caspase-1, active form of caspase-1 (p20), pro-IL-1β and mature IL-1β in the liver homogenates of WT and *Rip1^K45A/K45A^* mice which were fed with a normal diet (ND) or MCD diet. Data are expressed as mean ± SEM (n=6 or 12 per group). ns not significant, *\*p*<0.05, *\*\*p*<0.01, \*\*\**p*<0.001.

**Fig. 4.**
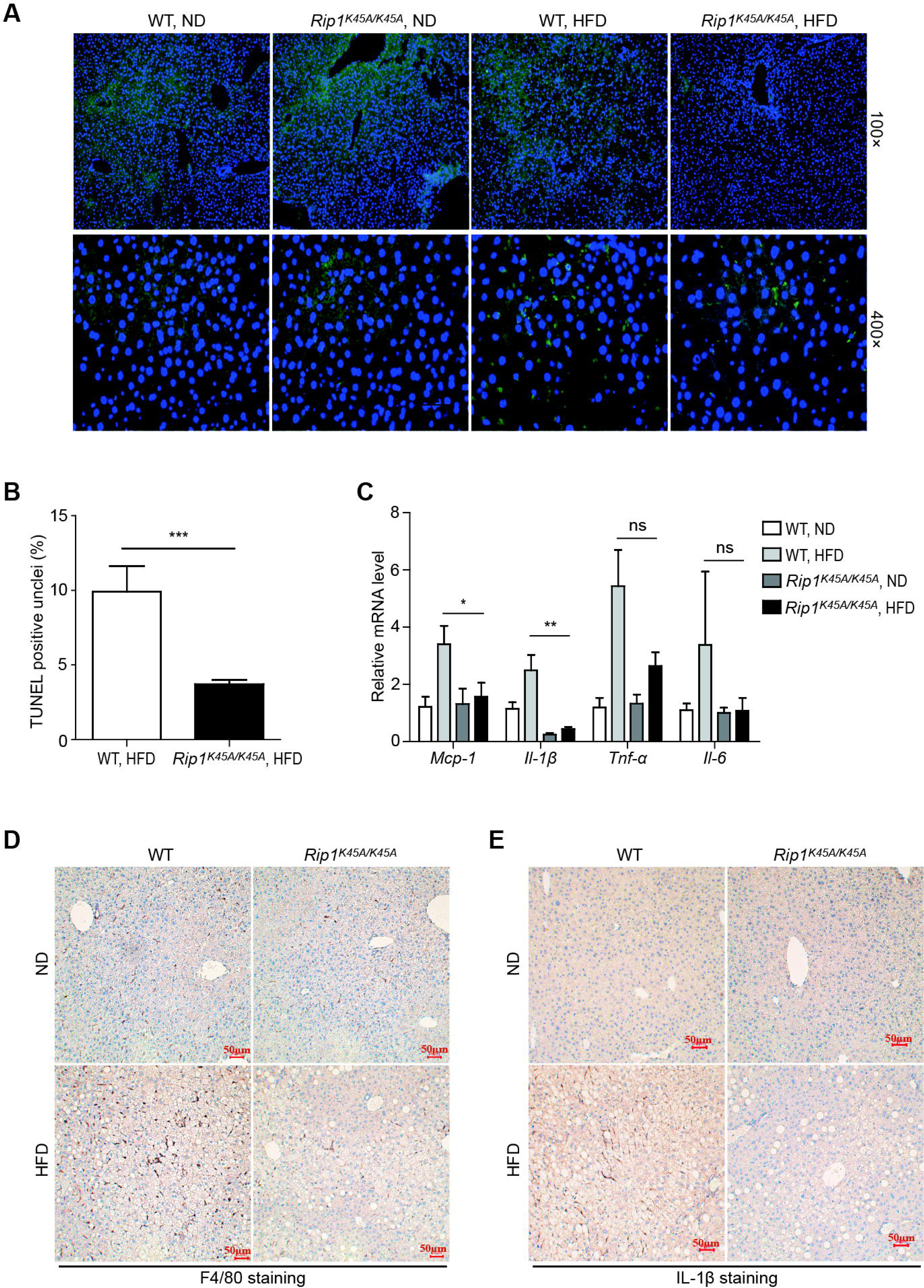
RIP kinase activity contributed to the hepatic cell death and inflammation in HFD-induced steatohepatitis. (A-B) Representative images and quantification of TUNEL-stained liver sections. (C) Hepatic mRNA expression of the inflammatory molecules *Mcp-1*, *IL-1β, Tnf-α* and *IL-6* in indicated groups. Representative images of (D) F4/80 staining and (E) IL-1β staining by immunohistochemistry analysis in WT and *Rip1^K45A/K45A^* mice fed with normal diet (ND) or HFD diet. Data are expressed as mean ± SEM (n=6 or 9 per group). ns not significant, *\*p*<0.05, *\*\*p*<0.01, \*\*\**p*<0.001.

### RIP1 kinase activity in hematopoietic-derived macrophages was critical for the development of steatohepatitis

Liver macrophages, including resident Kupffer cells and monocyte-derived macrophages, play critical roles in triggering hepatic inflammation in chronic liver diseases (22). Now it is widely accepted that the liver injury induced by lipotoxicity, stimulates the activation of Kupffer cell and recruitment of circulating bone marrow-derived monocytes into the liver, leading to sustained inflammation and fibrosis progression (23, 24). To further investigate the underlying mechanism and to distinguish the roles of hematopoietic or nonhematopoietic RIP1 kinase in NASH development, we used irradiation and bone marrow transplantation to create chimeric mice with RIP1 kinase inactivation only in hematopoietic cells (Fig. 5A). We first confirmed the efficiency of bone marrow transplantation (Fig. 5B). Following 5 weeks of MCD feeding, mice containing hematopoietic RIP1 K45A/K45A mutation exhibited extensively attenuated liver injury than WT➔ WT controls (Fig. 5C). It is noteworthy that the chimeric mice exhibited comparable and even more efficient protection against MCD-induced liver injury, steatosis and fibrosis than whole-body *Rip1^K45A/K45A^* mice, as illustrated by hepatic TG detection, H&E staining, Oil Red O staining, Sirius Red staining and α-SMA immunohistochemical staining of liver sections, and qPCR detection of fibrotic markers (*Col3a1*, *Tgf-β* and *α-sma*) (Fig. 5D-I). In addition, reduced expression of the inflammatory molecules (*Mcp-1, Il-1β, Tnf-α* and F4/80) confirmed that hepatic inflammation was inhibited in RIP1 KD chimeric mice (Fig. 5L&M). Moreover, percentage of TUNEL-positive cells was significantly decreased in the liver of KD➔WT chimeric mice (Fig. 5J&K). Taken together, these results suggest that the kinase activity of RIP1 in hematopoietic-derived macrophages determined its contribution to the inflammation, pathology and fibrosis development in steatohepatitis.

**Fig. 5.**
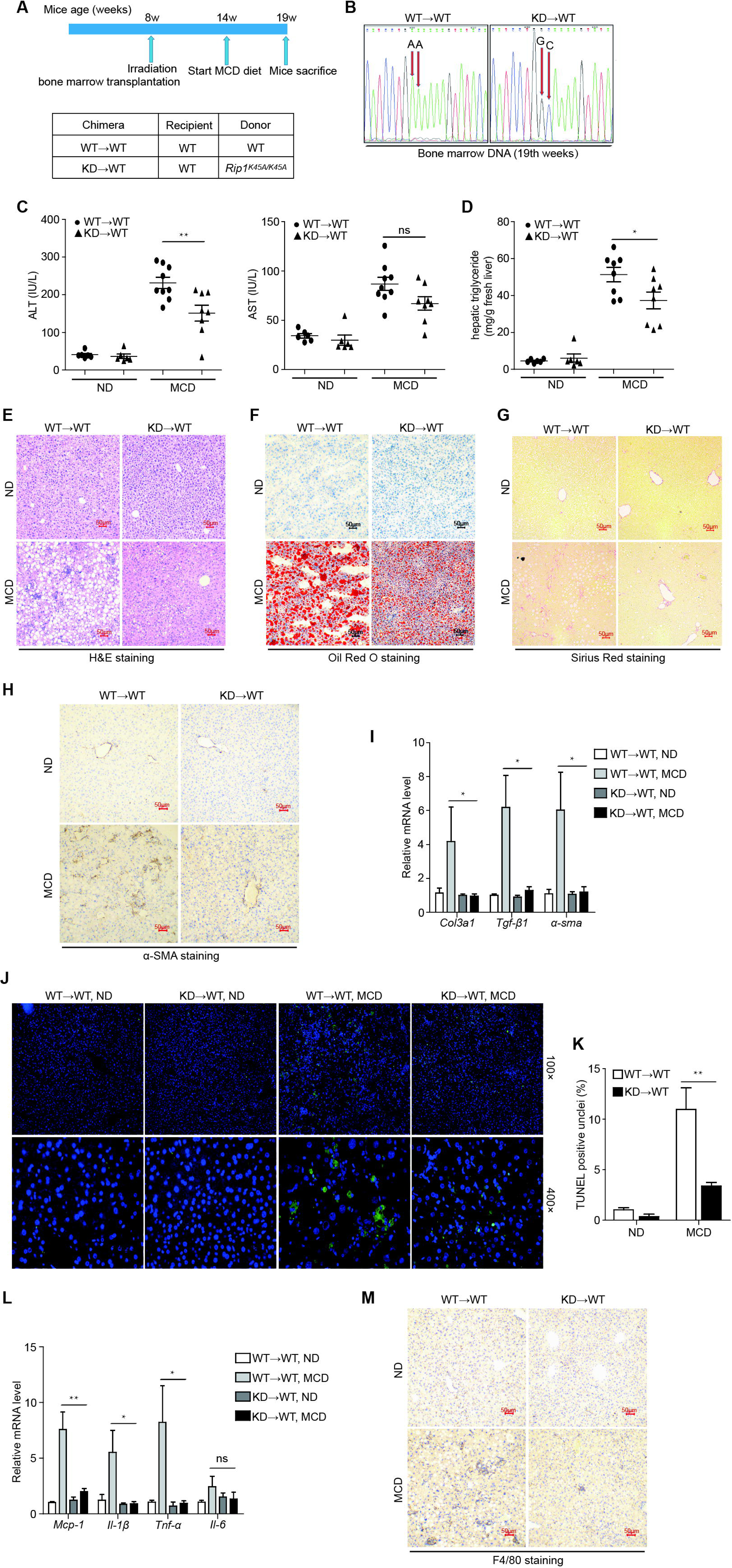
RIP1 kinase activity in hematopoietic-derived macrophages was crucial for the disease progression in steatohepatitis. (A) The experimental design is illustrated. (B) Reconstitution of donor bone marrow cells were verified by DNA sequencing. (C) Serum ALT and AST, (D) hepatic TG levels were analyzed respectively after 5-week MCD feeding. Liver sections were analyzed with (E) H&E staining, (F) Oil Red O staining, (G) Sirius Red staining and (H) α-SMA immunohistochemistry staining. Representative photographs of each group are presented. (I) Hepatic mRNA levels of the fibrosis markers *Tgf-β1*, *Col3a1* and *α-sma* in indicated groups. (J-K) Representative images and quantification of TUNEL-stained liver sections. (L) Hepatic mRNA expression of the inflammatory molecules *Mcp-1*, *IL-1β, Tnf-α* and *IL-6*. (M) Representative photographs of F4/80-stained liver sections detected by immunohistochemistry in different groups of mice fed with normal diet (ND) or MCD diet. Data are expressed as mean ± SEM (n=6 ∼ 9 per group). ns not significant, *\*p*<0.05, *\*\*p*<0.01, \*\*\**p*<0.001.

### RIP1 kinase was required for fatty acid-induced inflammasome activation and cell death

We postulated that the kinase activity of RIP1 contributes to the liver injury and inflammation induced by lipotoxicity in steatohepatitis. To model this *in vitro*, we used palmitic acid (PA) or oleic acid (OA), which are the dominant saturated or unsaturated free fatty acids (FFA) in human plasma, to treat bone marrow-derived macrophages (BMDMs) or AML12 mouse hepatocytes. Serum FFA especially PA levels are elevated in patients with NASH (25, 26) and have been shown to cooperate with gut-derived endotoxin (LPS) to contribute to the pathogenesis of NASH (27). Our results indicated that saturated fatty acid, palmitic acid (PA), other than unsaturated fatty acid OA, induced obvious cell death in BMDMs or AML12 mouse hepatocytes (Fig. S4B & S5), and the cell death was significantly rescued in *Rip1^K45A/K45A^* BMDMs or by RIP1 kinase inhibitor pretreatment (Fig. 6A&B, S5). In agreement with previous studies (27, 28), PA also induced inflammasome activation, which caused the secretion of IL-1β in a dose-dependent manner in LPS-primed BMDMs (Fig. 6C). The kinase activity of RIP1 was not required for the “signal 1” of inflammasome activation, as the TNF-α production which is controlled by NF-κB pathway (“signal 1”) was not influenced by RIP1 kinase inactivation or inhibition (Fig. S3). In contrast, cleavage of caspase-1 and IL-1β release were markedly reduced in *Rip1^K45A/K45A^* BMDMs or Nec-1s-pretreated BMDMs (Fig. 6E-H), suggesting that RIP1 kinase contributed to the inflammasome activation mainly through mediating “signal 2” and this result is in line with previous findings (16). In addition, PA treatment also induced the phosphorylation (S166) of RIP1 in BMDMs (Fig. 6I), and this was consistent with our *in vivo* results (Fig. 1A), suggesting that RIP1 kinase can be activated by either saturated fatty acid *in vitro* or lipotoxicity *in vivo*. Taken together, these results suggest that RIP1 kinase activity played a key role in mediating the cell death and inflammasome activation induced by saturated fatty acid PA, which has been recognized as an important trigger for NASH.

**Fig. 6.**
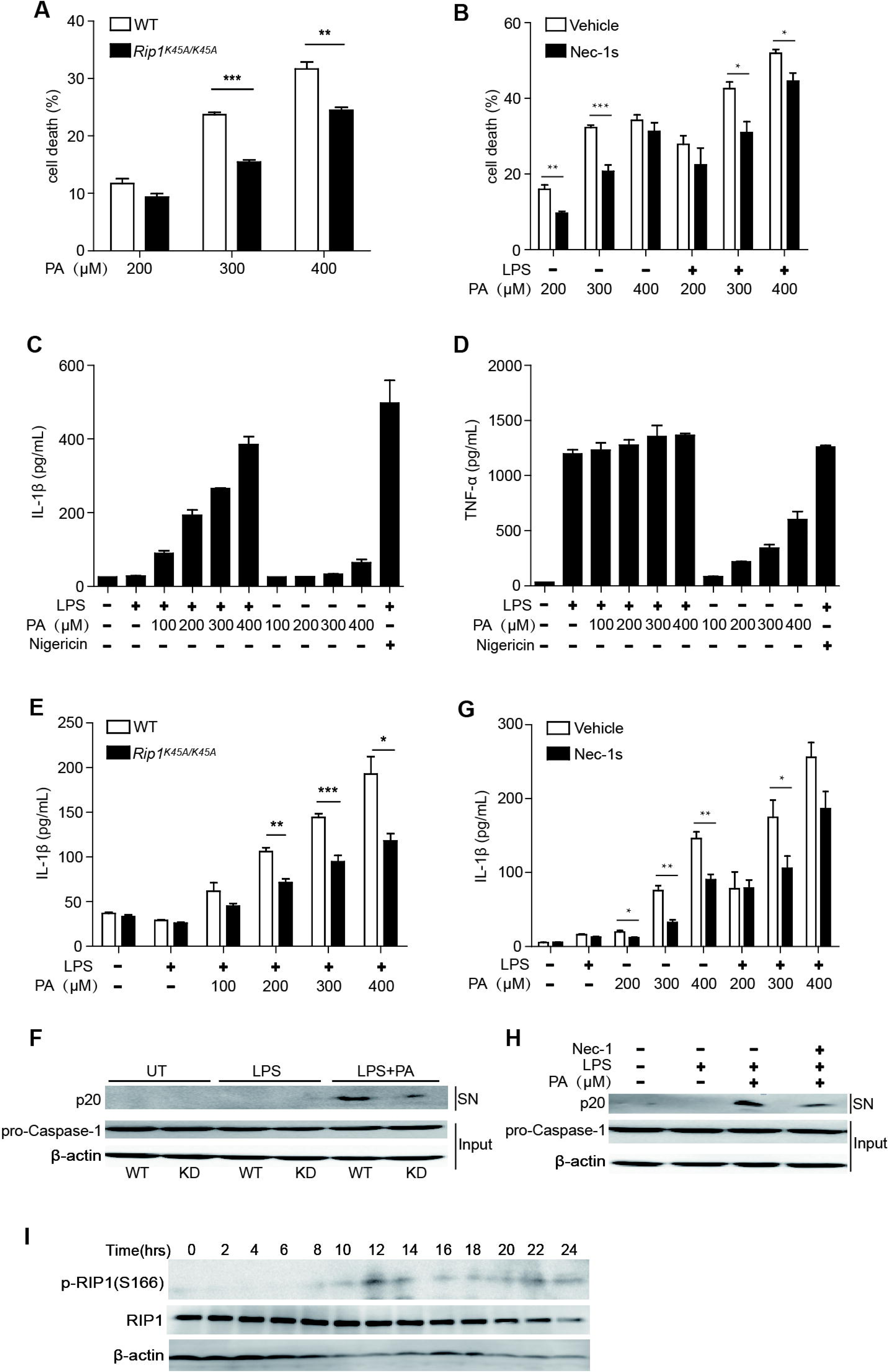
RIP1 kinase contributed to saturated fatty acid-induced cell death and inflammasome activation in macrophages. (A) Bone marrow-derived macrophages (BMDMs) isolated from WT or *Rip1^K45A/K45A^* mice were treated with different concentrations of PA as indicated for 24 h and then the cell death was evaluated by LDH release assay. (B) With or without pretreatment by RIP1 kinase inhibitor Nec-1s (40 μM) for 1 hour, WT BMDMs were treated as indicated and the cell death rate was measured. (C-D) WT BMDMs were stimulated by different doses of PA with or without LPS priming. The cytokine IL-1β and TNF-α in supernatant were measured by ELISA. (E-F) WT or *Rip1^K45A/K45A^* BMDMs were primed with LPS and then treated with different concentrations of PA. (G-H) With or without Nec-1s pretreatment, BMDMs were treated with PA or LPS plus PA. (E&G)Then the secretion of IL-1β cytokine in supernatant was detected by ELISA. (F&H) The cleaved caspase-1 (p20) in supernatants (SN) or pro-caspase-1 and β-actin in cell lysates (Input) were analyzed using immunoblot. (I) BMDMs were treated with PA (400 μM) for indicated time. The S166 phosphorylation and expression of RIP1 were detected by immunoblot using respective antibodies. Figures are representative of three independent experiments. Data are expressed as mean ± SEM. ns not significant, *\*p*<0.05, *\*\*p*<0.01, \*\*\**p*<0.001.

### Expression and phosphorylation of RIP1 was increased in liver tissues of NAFLD or NASH patients

The above results indicated that RIP1 kinase is important in experimental steatohepatitis. We then wondered whether it was also involved in human NASH development. First, analysis of the public GEO database suggested that the expression of RIP1 was robustly elevated in liver tissues of NASH or NAFLD patients, similar as in murine models of NASH (Fig. 7A and Fig. S6A-C). We also examined whether RIP1 was phosphorylated and activated in the liver tissues of NAFLD or NASH patients. The patients’ information was summarized in Table 1. As determined by immunofluorescence staining analysis, the phosphorylation of RIP1 (S166) was obviously triggered in patients with NASH, but not in healthy controls (Fig. 7E). The increased expression and phosphorylation of RIP1 was correlated with the steatosis, liver injury and fibrosis in the NASH livers (Fig. 7B-D). These results suggested that in NAFLD or NASH patients, both the expression and activation of RIP1 kinase was stimulated and increased.

**Table 1.**

| Diagnosis | Number | Age | Sex | ALT<br>(U/L) | AST<br>(U/L) | TG<br>(mmol/L) | TC<br>(mmol/L) | HDL<br>(mmol/L) | LDL<br>(mmol/L) | Glucose<br>(mmol/L) |
| --- | --- | --- | --- | --- | --- | --- | --- | --- | --- | --- |
| normal | 1 | 58 | male | 26.5 | 25.9 | 0.98 | 4.65 | 1.48 | 2.87 | 4.94 |
|  | 2 | 36 | male | 36.7 | 22 | 1.54 | 5.43 | 1.55 | 3.31 | 4.11 |
|  | 3 | 47 | male | 20.5 | 16.1 | 1.66 | 3.98 | 0.94 | 2.45 | 5.89 |
|  | 4 | 39 | male | 11.9 | 17.4 | 1.11 | 4.33 | 1.22 | 2.67 | 4.66 |
|  | 5 | 37 | female | 11.5 | 18.6 | 1.1 | 4.87 | 1.18 | 3.05 | 4.82 |
|  | 6 | 32 | female | 16.7 | 17.9 | 0.59 | 4.8 | 1.65 | 2.83 | 4.75 |
|  | 7 | 31 | female | 9.6 | 13.6 | 0.63 | 4 | 1.98 | 1.89 | 4.42 |
|  | 8 | 23 | female | 12.3 | 20 | 0.71 | 4.18 | 1.69 | 2.26 | 4.49 |
| NAFLD | 9 | 38 | female | 40.5 | 33.3 | 0.93 | 4.58 | 1.53 | 2.5 | 6.17 |
|  | 10 | 64 | female | 72 | 43.1 | 1.33 | 5.82 | 1.24 | 3.83 | 4.68 |
|  | 11 | 42 | female | 25.4 | 24.5 | 1.99 | 4.14 | 0.85 | 2.46 | 6.25 |
|  | 12 | 59 | female | 21 | 19 | 1.24 | 2.84 | 1.17 | 1.19 | 5.12 |
|  | 13 | 16 | female | 40.5 | 24.7 | 0.47 | 2.75 | 0.9 | 1.57 | 4.26 |
|  | 14 | 36 | female | 147.8 | 90 | 1.99 | 5.06 | 0.96 | 3.12 | 10.99 |
|  | 15 | 34 | female | 33.8 | 22 | 0.9 | 4.79 | 1.26 | 3.12 | 4.69 |
|  | 16 | 54 | female | 23.3 | 24.8 | 2.7 | 3.81 | 0.73 | 2.16 | 5.01 |
|  | 17 | 35 | female | 30.4 | 17.6 | 0.79 | 3.69 | 1.13 | 2.1 | 6.02 |
|  | 18 | 37 | male | 34.7 | 20.5 | 1.5 | 3.74 | 0.79 | 2.33 | 4.61 |
|  | 19 | 70 | male | 11.4 | 14.1 | 0.98 | 4.43 | 1.14 | 2.81 | 4.2 |
|  | 20 | 57 | male | 34.6 | 20.2 | 0.72 | 3.7 | 1.21 | 2.04 | 4.68 |
|  | 21 | 35 | male | 64.5 | 36.2 | 2.95 | 4.91 | 0.73 | 3.25 | 6.72 |
|  | 22 | 32 | male | 29.2 | 23.2 | 1.63 | 3.94 | 0.88 | 2.36 | 5.07 |
|  | 23 | 25 | male | 78.4 | 74.5 | 1.71 | 4.83 | 0.89 | 3.12 | 5.17 |
|  | 24 | 14 | male | 20.9 | 13.7 | 1.03 | 5.07 | 1.11 | 3.43 | 4.53 |
|  | 25 | 20 | male | 97.7 | 41.5 | 1.25 | 3.75 | 0.86 | 2.33 | 4.66 |
|  | 26 | 31 | male | 152.4 | 64.1 | 2.06 | 6.1 | 1.25 | 4.21 | 5 |
|  | 27 | 49 | male | 94 | 45.7 | 1.3 | 4.47 | 0.98 | 2.84 | 5.34 |
|  | 28 | 19 | male | 30 | 27.1 | 1.07 | 5.47 | 1.14 | 3.61 | 4.73 |
|  | 29 | 26 | male | 127.4 | 87 | 2.71 | 5.39 | 0.84 | 3.45 | 6.91 |
|  | 30 | 26 | male | 169.9 | 61.3 | 1.71 | 4.83 | 0.99 | 3.26 | 4.63 |

**Fig. 7.**
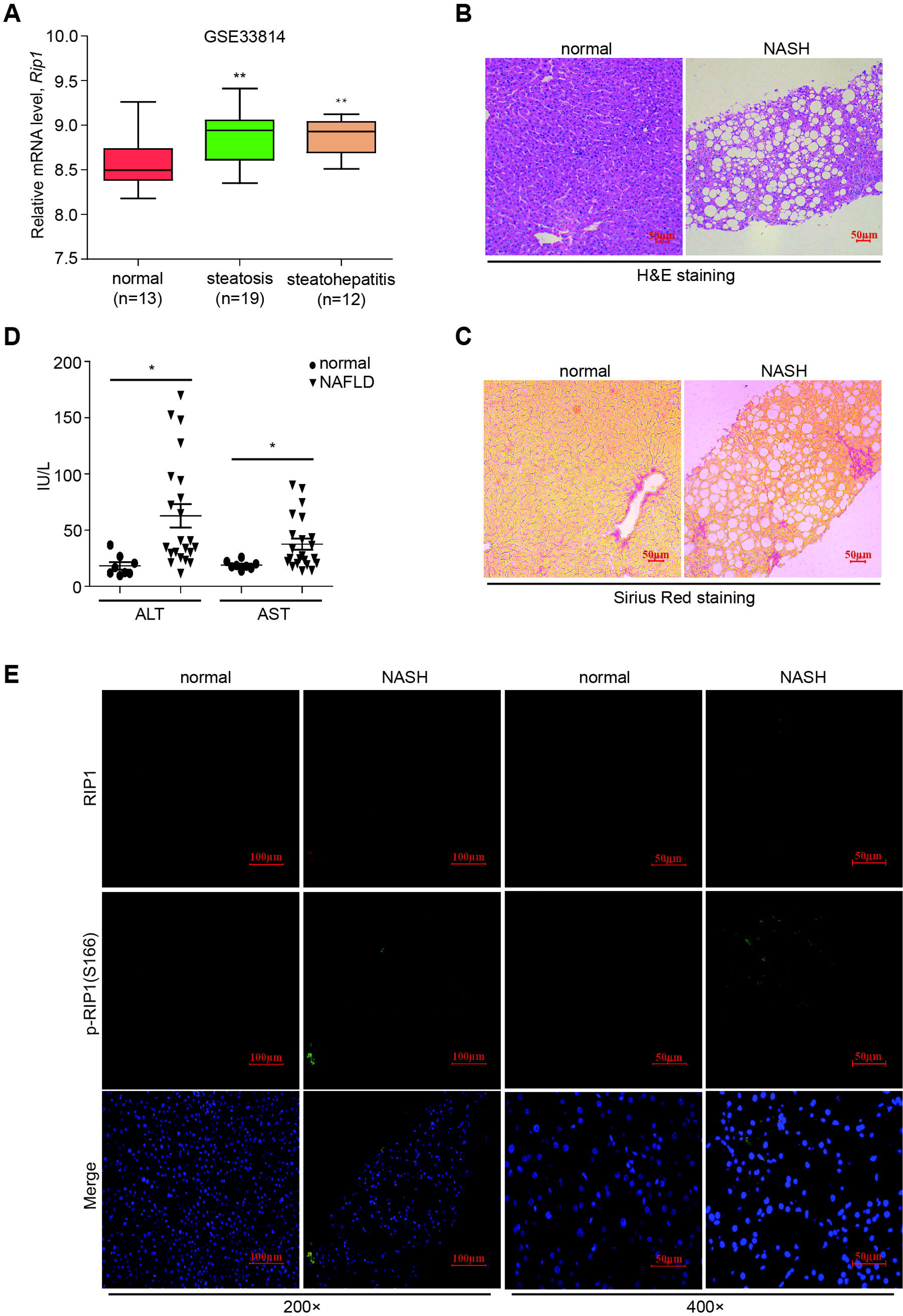
RIP1 expression and activation were increased in liver tissues of NAFLD or NASH patients. (A) Analysis of mRNA levels of RIP1 using published datasets to compare RIP1 expression between healthy controls and NAFLD or NASH patients. (B-C) Representative photographs of (B) H&E staining and (C) Sirius Red staining of liver sections from normal persons or NASH patients. (D) The serum ALT and AST values of the collected NAFLD or NASH patients and the healthy controls. (E) Representative images of liver tissue sections from control or NASH patients analyzed for the expression and phosphorylation of RIP1 by immunofluorescence staining. Data are expressed as mean ± SEM. ns not significant, *\*p*<0.05, *\*\*p*<0.01, \*\*\**p*<0.001.

## Discussion

The most important finding in the present study is that we for the first time demonstrated the critical contribution of the kinase activity of RIP1 in hematopoietic-derived macrophages to the pathology of steatohepatitis. Hepatic RIP1 expression and kinase activation (phosphorylation) increases in both experimental and clinical NASH. Our results not only provide compelling evidence to support earlier work by Majdi et al. for a critical role of RIP1 kinase in pathogenesis of NASH (29), but also further suggest a special and key function of macrophages to the NASH development via RIP1 kinase activity. It is recently reported by Tao et al. on *Nature* journal that different human cell types exhibit opposite phenotypes regarding RIP1 kinase-mediated cell death and inflammation (30). Combined with our results, it is therefore suggested that when developing pharmaceutical strategy targeting RIP1 kinase, it would be more safe and beneficial to target RIP1 kinase in specialized cell type(s), for example, in macrophages and monocytes.

In our study, we have established two entirely different murine models of steatohepatitis, both of which have been extensively used in NASH research but emphasizing different characteristics and mechanisms regarding steatohepatitis (31). MCD model exhibits hallmarks of NASH pathology, including steatosis, significant lobular inflammation and liver fibrosis, as often observed in human NASH. But mice in MCD model lose body weight and this does not mimic human NASH. In contrast, HFD model is an overnutrition model in which NASH accompanies the development of obesity and related metabolic syndromes, including insulin resistance and glucose intolerance, which are invariable characteristics in human NASH (31, 32). RIP1 kinase-dead (*Rip1^K45A/K45A^*) mice exhibited significantly attenuated liver injury, inflammation as well as fibrosis in both models, suggesting that the involvement of RIP1 kinase in NASH pathogenesis is not limited to specific model, but might generally apply to models induced by different factors or conditions.

Hepatocellular death represents a central feature in chronic liver disease, contributing to the recruitment of immune cells, activation of hepatic stellate cells and the development of liver fibrosis, cirrhosis and even cancer. In terms of the cell death mode in steatohepatitis, earlier studies suggested that hepatocytes died mainly by apoptosis, a classical programmed and non-inflammatory mode of cell death (33). Cleaved caspase-3 and other apoptosis markers like CK18 were detected in samples from NASH patients (34, 35). Genetic or pharmaceutical inhibition of caspases has been shown to attenuate liver injury and liver fibrosis in experimental NASH models or in clinical trials (36). However, along with the advances in knowledge regarding different forms of cell death and the relevant molecular mechanisms, growing evidences suggest that the mode of hepatocellular death during steatohepatitis is not limited to apoptosis, and another programmed and inflammatory cell death mode, necroptosis, is also involved (8). The critical regulator and executioner of necroptosis, RIP3 and MLKL, have been demonstrated to play roles in NASH development, despite that there were controversies regarding the role of RIP3 in NASH (37–39). As an important regulator of both apoptosis and necroptosis, RIP1 has been intensively studied in different liver diseases, including acute liver failure and hepatocellular carcinoma (40–42). In Concanavalin A (ConA)-induced liver injury model, the mice with RIP1 deficiency in liver parenchymal cells (*Ripk1^LPC-KO^*) showed opposite phenotypes with the RIP1 kinase-dead (*Rip1^K45A/K45A^*) mice, suggesting that the kinase activity of RIP1 play distinct roles than its scaffolding function in liver injury (43). Using specific RIP1 kinase-dead mice, our study provided direct genetic evidence that the kinase activity of RIP1 is important for promoting the pathological progression in NASH. But regarding the underlying mechanisms, in *Rip1^K45A/K45A^* mice, either MCD or HFD feeding-induced cell death (TUNEL-positive cells) in liver tissue were significantly reduced (around 50% reduction), other than completely inhibited. These *in vivo* results were in agreement with *in vitro* model, that saturated fatty acid PA-induced cell death was significantly decreased but not completely inhibited by RIP1 kinase inhibitor or in BMDMs from *Rip1^K45A/K45A^* mice. These results suggest that in addition to apoptosis and necroptosis mediated by RIP1 kinase, other cell death modes might be also involved. Xu et al. reported that the critical pyroptosis executor Gasdermin D (GSDMD) plays a key role in NASH in both humans and mice, suggesting that another inflammatory cell death mode, pyroptosis, at least partially contributes (44). During the progression of steatohepatitis, lipotoxitiy caused by steatosis, and other risk factors including oxidative stress, ER stress and mitochondria disfunction, result in a complex microenvironment in liver and lead to the induction of different cell death modes via different mechanisms. Hence, the benefit will be limited for just targeting one specialized cell death machinery, and the previous studies using either MLKL^-/-^, RIP3^-/-^, GSDMD^-/-^ mice or pan-caspase inhibition which all exhibited only partial protection against steatohepatitis also supported this opinion (38, 39, 44).

During the development of steatohepatitis, chronic hepatic inflammation also plays an important role. Upon inflammation induction, massive monocytes will be recruited to the sites of liver injury. Monocyte-derived macrophages thereby turn to be the dominant population among hepatic macrophages, which further aggravate hepatic inflammation, activate HSCs and prolong the survival of activated HSCs, thus leading to the progression of liver fibrosis (23, 24). Therefore, macrophage has been proposed as a potential therapeutic target for developing novel strategies to treat chronic liver diseases (22). Krenkel et al. showed that inhibition of monocyte recruitment could efficiently reduce steatohepatitis and fibrosis (24). In line with these findings, our study indicated that the kinase activity of RIP1 in monocyte-derived macrophage plays a critical role in promoting the fibrosis in NASH. The chimeric mice with kinase-inactive RIP1 only in hematopoietic cells exhibited similar protection against NASH as whole-body *Rip1^K45A/K45A^* mice. For the underlying mechanism, accumulating evidence suggest that NLRP3 inflammasome activation and the production of pro-inflammatory cytokine IL-1β by hepatic macrophages play crucial roles in liver fibrosis (45, 46). In agreement, our *in vitro* and *in vivo* results demonstrated that the kinase activity of RIP1 contributed to the pathogenesis and fibrosis in steatohepatitis at least partially through mediating the inflammasome activation, caspase-1 cleavage and IL-1β production in liver macrophages.

In our study, we confirmed the clinical relevance of RIP1 kinase activation (evaluated by RIP1 phosphorylation) and expression in a cohort of NASH patients. We also searched in open database and analyzed the expression of RIP1 in different published studies, and found that the expression level of RIP1 correlated with the disease progression in NAFLD or NASH.

In summary, our study provided the direct genetic evidence that RIP1 kinase plays a key role in the development of NASH using two different murine models, supporting the potential application of RIP1 kinase inhibitor in treating steatohepatitis. We offer an alternative explanation regarding the mechanism that RIP1 kinase contributes to the pathogenesis of NASH via mediating cell death and inflammation, especially inflammasome activation. Moreover, our results also suggest that pharmaceutical targeting RIP1 kinase activity in specialized cell type such as monocyte-derived macrophage might be more beneficial.

## Materials and Methods

### Reagents

Necrostatin-1 (Nec-1) and Necrostatin-1s (Nec-1s) were purchased from Enzo Lifesciences (Switzerland). Anti-Caspase-1 (p20) (mouse) mAb (Casper-1) (AG-20B-0042-C100) was from Adipogen (Germany). Anti-RIP1 (mouse) mAb was from BD Biosciences (America). Phospho-RIP (Ser166) ((D8I3A) Rabbit mAb #44590)/(Antibody (Rodent Specific) #31122), α-Smooth Muscle Actin ((D4K9N) XP® Rabbit mAb #19245), F4/80 ((D2S9R) XP® Rabbit mAb #70076) and IL-1β ((3A6) Mouse mAb #12242) were from Cell Signaling Technology. Lipopolysaccharides (LPS) was from Sigma-Aldrich. Nigericin (481990) was from Merck-Millipore (Germany). Insulin, Transferrin, Selenium Solution (ITS-G), 100X (41400045) and Penicillin-Streptomycin, Liquid (15140122) were from GIBCO (America). Hifair® Ⅱ 1st Strand cDNA Synthesis Kit (11119ES60) was from Yeasen (Shanghai, China). KAPA SYBR^®^ Fast (KK4601) was from KAPA BIOSYSTEMS (America). IL-1β ELISA kit (Catalog # 88-7013-22) and TNFα ELISA kit (Catalog # 88-7324-88) were from Invitrogen (America). LDH Cytotoxicity Assay Kit (C00170) was from Beyotime (Shanghai, China). Alanine transaminase (ALT) detection kit (C009-2-1), aspartate transaminase (AST) detection kit (C010-2-1), Triglyceride (TG) detection kit (A110-1-1) or (BC0625), and cholesterol (TC) detection kit (A111-1-1) were all purchased from Nanjing Jiancheng Bioengineering Institute (Nanjing, China) or Solarbio Science & Technology (Beijing, China).

### Mouse models

C57BL/6 mice were purchased from Model Animal Research Center of Nanjing University (Nanjing, China). *Rip1^K45A/K45A^* mice on a C57BL/6 background as previously described (18) were kindly provided by Dr. Haibing Zhang (Chinese Academy of Sciences, Shanghai, China). All animals were maintained under standard laboratory conditions, with free access to food and water. All animal experiments were performed under protocols approved by the Animal Care and Use Committee at Nanjing University of Science & Technology.

Eight-week-old male *Rip1^K45A/K45A^* mice (C57BL/6 background) and male wild-type C57BL/6 controls were randomly grouped and fed either with methionine-choline deficient diet (MCD, TP3006) or control diet (Trophic Animal Feed, China) for five weeks to induce steatohepatitis. Separate groups of *Rip1^K45A/K45A^* and WT mice were fed either with a high-fat diet (HFD, 60.9% fat, 18.3% protein, 20.8% carbohydrate, TP23520; Trophic Animal Feed, China) or control diet (Trophic Animal Feed, China) for 24 weeks to trigger HFD-induced steatohepatitis. During the HFD feeding period, body weight gain, food and water consumption, fasting blood glucose were monitored or detected. At the end of HFD feeding, metabolic parameters including O_2_ consumption, CO_2_ production, respiratory exchange ratio (RER) as well as heat production were measured by metabolic cage detection (Comprehensive Lab Animal Monitoring System). Glucose tolerance test (GTT) was also performed as previously described (47).

For bone marrow transplantation, the recipient wild-type C57BL/6 mice were lethally irradiated (9 Gy) and then injected through tail veins with bone marrow cells (5 × 10^6^ cells) harvested from wild-type or *Rip1^K45A/K45A^* donor mice. 6 weeks after irradiation and bone marrow transplantation (BMT), the chimeric mice were fed with either MCD diet or normal diet for another 5 weeks to induce NASH. Bone marrow cells and blood cells of the chimeric mice were also collected and analyzed by DNA sequencing to verify the bone marrow reconstitution efficiency.

### Human samples

Human liver tissue samples and corresponding clinical information were provided by Nanjing Drum Tower Hospital (Nanjing, China). Paraffin-embedded human liver tissue sections were analyzed by H&E staining or Sirius Red staining. Expression and phosphorylation (S166) of RIP1 were detected by immunofluorescence staining using specific antibodies. All individuals gave written informed consent before joining this study. All research procedures were approved by the Clinical Research Ethics Committee of the Nanjing Drum Tower Hospital.

### Cell culture and treatments

Mouse bone marrow-derived macrophages (BMDMs) were derived by maturing bone marrow cells from adult wild-type or *Rip1^K45A/K45A^* mice in the presence of M-CSF containing supernatant from L929 cells as previously described (48). AML-12 mouse hepatocytes, which were purchased from the Cell Bank of the Chinese Academy of Sciences (Shanghai, China), were cultured in DMEM (GIBCO) with 10% heat-inactivated FBS, 1% penicillin-streptomycin, 1% Insulin, Transferrin, Selenium Solution (100X) and 4.6 μL dexamethasone (5 mg/mL).

Some cells were primed with LPS (100 ng/mL) for 3 hours and then cells were treated with different concentrations of saturated fatty acid palmitic acid (PA) or unsaturated fatty acid oleic acid (OA) for 24 hours. Then cell supernatant was harvested and the cytokine IL-1β (Catalog # 88-7013-22, Invitrogen) and TNFα (Catalog#88-7324-88, Invitrogen) in cell supernatants were measured using Enzyme-linked Immunosorbent Assay (ELISA) kits according to the manufacturer’s instructions. Cell lysates were also harvested and analyzed by immunoblotting. In addition, cell death rate was quantified by detecting the lactate dehydrogenase (LDH) release using LDH Cytotoxicity Assay Kit (C00170; Beyotime, China) following the manufacturer’s instruction

### Immunoblotting

Mouse liver samples, BMDMs and AML-12 cells were lysed in RIPA cell lysis buffer with protease inhibitor (P1005; Beyotime, China) and phosphatase inhibitor (P1081; Beyotime, China). Equal amounts of proteins were separated by SDS-PAGE (Acryl/Bis 30% solution (29:1) (B546017; Sangon Biotech, China), transferred to 0.22 μm PVDF membranes, blocked with 5% skim milk powder in TBS-T (0.1% TWEEN-20). The membranes were incubated with primary antibodies at 4°C overnight, washed with TBS-T and incubated with secondary antibodies at room temperature for 1-2 hour. Chemiluminescent Substrate System from KPL was utilized for final detection.

### TUNEL assay

Apoptotic cells in livers were measured by TUNEL Apoptosis Assay Kits. Paraffin-embedded liver tissue sections were pretreated by proteinase K for 20 min at 25°C. Then tissue sections were washed 3 times with PBS and then incubated in the mixture of reaction buffer with TdT enzyme in dark for 60 min at 37°C. Meanwhile the cell nucleuses were stained with DAPI. The tissue sections were observed and photographed using the fluorescence microscope (NIKON ECLIPSE 80i). The apoptotic cell rate was calculated with the formula: (TUNEL positive cell number / total area cell number) ×100%.

### mRNA isolation and qPCR analysis

TRIzol (Invitrogen) reagent was added to cells or tissues and total RNA was extracted according to the manufacturer’s instructions. Next cDNA was synthesized using Hifair® II 1st Strand cDNA Synthesis Kit. PCR reactions were performed on the ABI 7300 real-time PCR system using KAPA SYBR^®^ Fast. GAPDH mRNA was used as an internal control to normalize mRNA expression. The sequences of primers for qPCR were as follows:

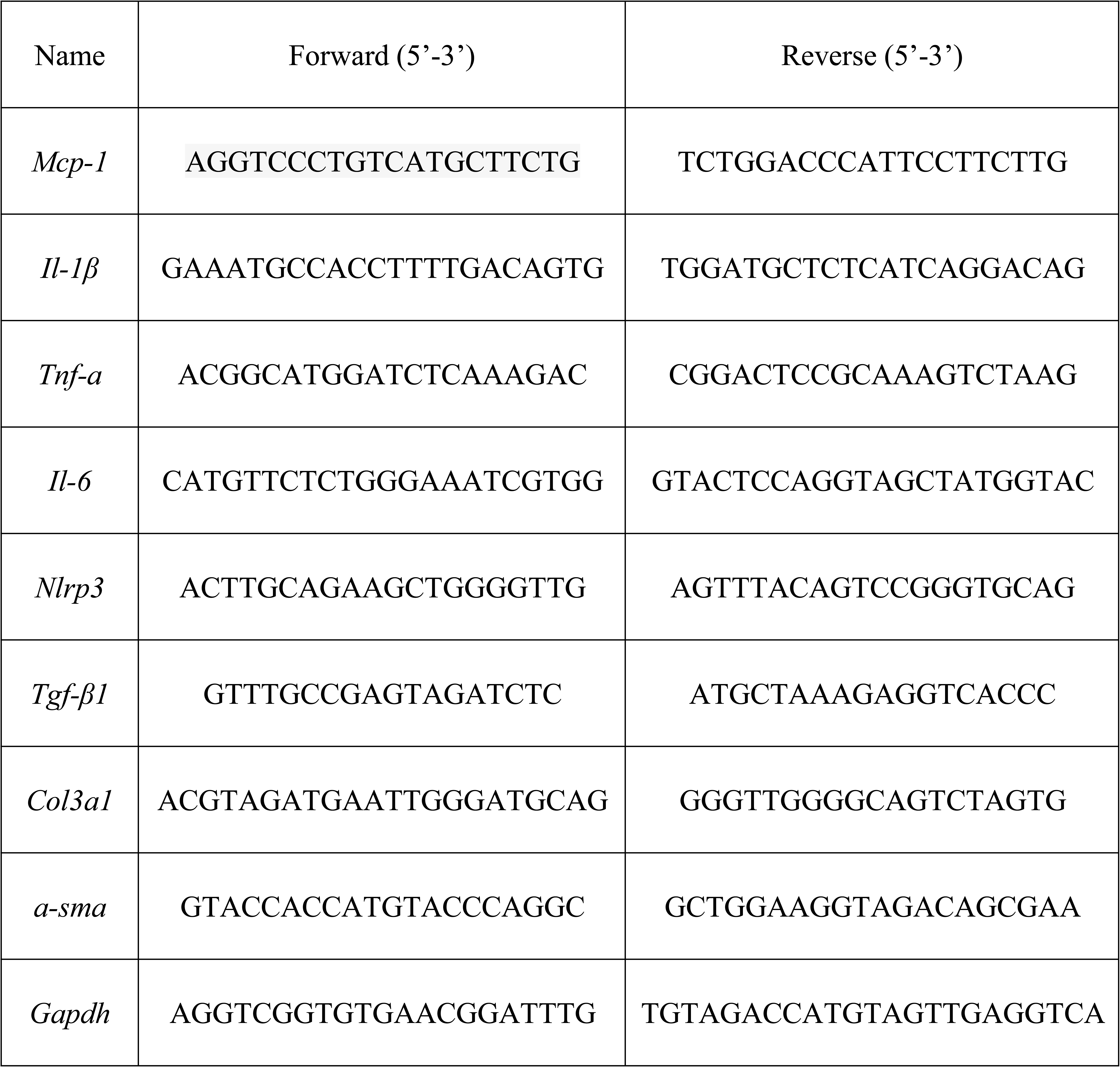

### Histological analysis

Mouse liver tissues were fixed in 4% paraformaldehyde and then embedded in paraffin and sectioned. Paraffin-embedded liver sections were stained with hematoxylin and eosin (H&E) to evaluate the gross morphology or with Sirius Red to evaluate liver fibrosis. At the same time, frozen sections of liver tissues were stained by Oil Red O working solution to determine the steatosis in liver tissues.

### Immunohistochemistry and immunofluorescence analysis

Paraffin-embedded liver sections were firstly deparaffinized and then boiled in citrate buffer (pH 6.0) for antigen retrieval, followed by hydrogen peroxide/PBS blocking of endogenous peroxidase. Slides were pre-blocked and then incubated with primary antibodies at 4 °C overnight. Primary antibodies, including the anti-F4/80, anti-ɑ-SMA and anti-IL-1β antibodies were used respectively. The slides were then washed with PBS and incubated with secondary antibodies at 37°C for 50 min, and then stained with DAB substrate after 20 min with streptavidin-HRP. The cell nucleuses were stained with hematoxylin. For immunofluorescence, after being incubated with primary antibodies (anti-RIP1 and anti-Phospho-RIP (Ser166)) at 4°C overnight, slides were washed with PBS for 3 times and then incubated with fluorescent secondary antibody in dark for 60 min. The cell nucleuses were stained by DAPI. At last, the slides were observed and photographed using the fluorescence microscope (NIKON ECLIPSE 80i).

### Biochemical analysis

Serum levels of alanine transaminase (ALT), aspartate transaminase (AST), Triglyceride (TG) and cholesterol (TC) were analyzed by respective detection kits according to the manufactures’ protocols.

### Sequencing Data analysis

Several datasets which was published in public database were utilized for human or murine *Rip1* expression analyses, including *GSE33814* (49), *GSE63067* (50), *GSE46300* (51), *GSE35961* (52), all of which can be downloaded from GEO database.

### Statistical analysis

All the data are expressed as the mean ± standard error of the mean (SEM). The statistical analysis of the results was performed using GraphPad Prism® 7.01 software (San Diego, USA). Unpaired Student’s *t* test or one-way ANOVA (for more than 2 groups) analysis were used to calculate the differences in mean values. p<0.05 was considered to be a statistically significant difference.

## Acknowledgments

This study was supported by the National Natural Science Foundation of China under Grant 31970897, 21677076 and 31500698, Outstanding Youth Foundation of Jiangsu Province (BK20190069), the Natural Science Foundation of Jiangsu Province of China under Grant BK20150772, the Fundamental Research Funds for the Central Universities No.30919011102, Qing Lan Project of Jiangsu Province.

---

**Supplementary Figure 1.**
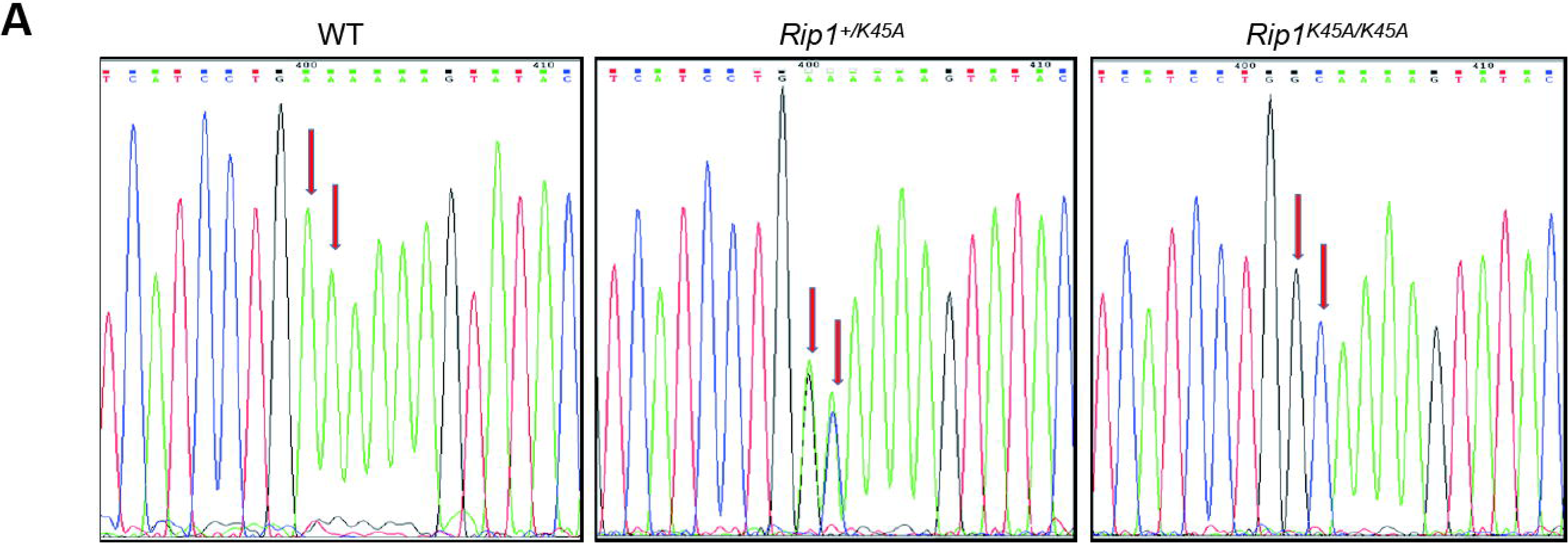
The genotype of *Rip1^K45A/K45A^* mice was confirmed by genomic DNA sequencing.

**Supplementary Figure 2.**
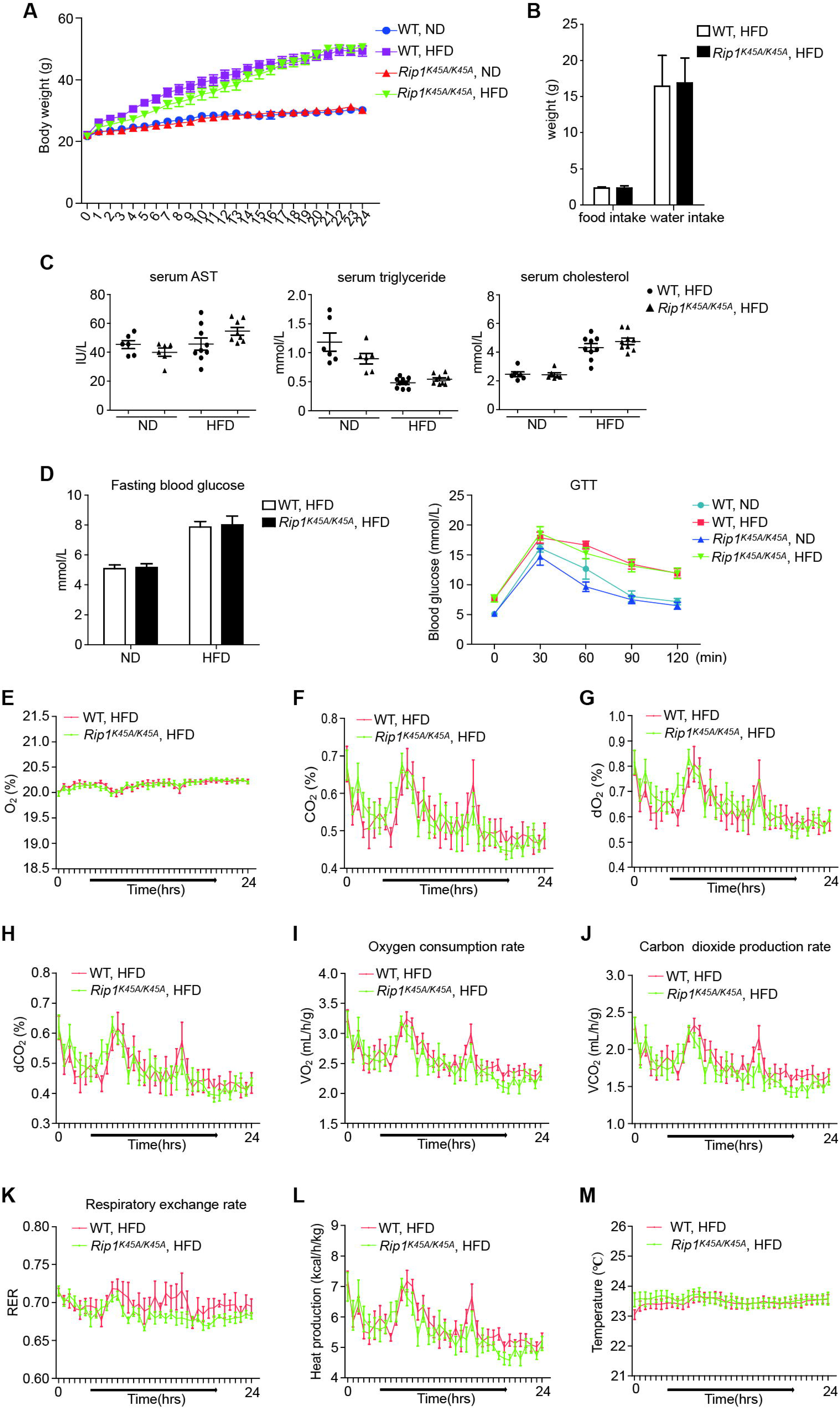
WT and *Rip1^K45A/K45A^* mice exhibited no obvious difference in body weight gain (A), food/water intake (B), serum levels of AST, TG and cholesterol (C), fasting blood glucose and glucose tolerance (D), oxygen consumption and CO_2_ production (E-J), respiratory exchange ratio (K), and physical activities (L&M) after ND or HFD feeding. Data are expressed as mean ± SEM (n=6 or 9 per group). ns not significant, *\*p*<0.05, *\*\*p*<0.01, \*\*\**p*<0.001.

**Supplementary Figure 3.**
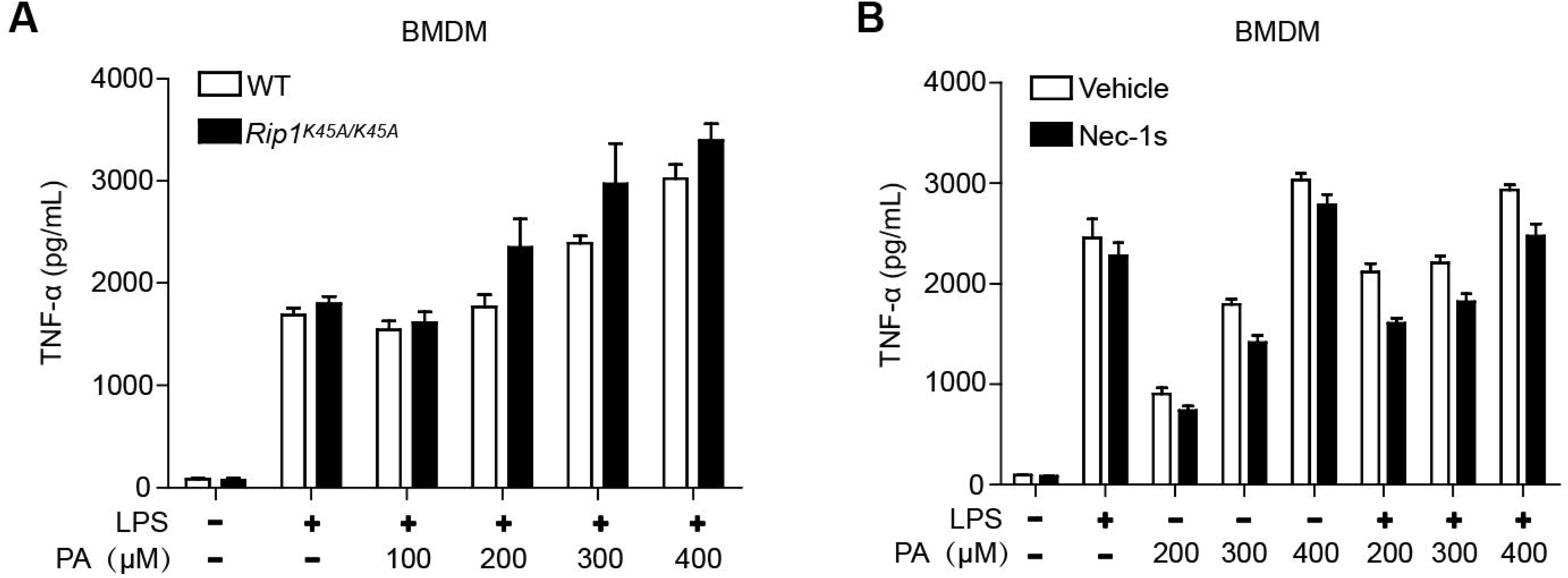
RIP1 kinase inactivation or inhibition did not affect the “signal 1” for inflammasome activation as characterized by TNFα production. (A) BMDMs from WT or *Rip1^K45A/K45A^* mice were stimulated by LPS plus PA as indicated for 24 hours. (B) WT BMDMs were pretreated with Nec-1s or vehicle control and then stimulated by PA or LPS plus PA for 24 hours. Secretion of TNFα in supernatant was measured by ELISA. Data are expressed as mean ± SEM. ns not significant, *\*p*<0.05, *\*\*p*<0.01, \*\*\**p*<0.001.

**Supplementary Figure 4.**
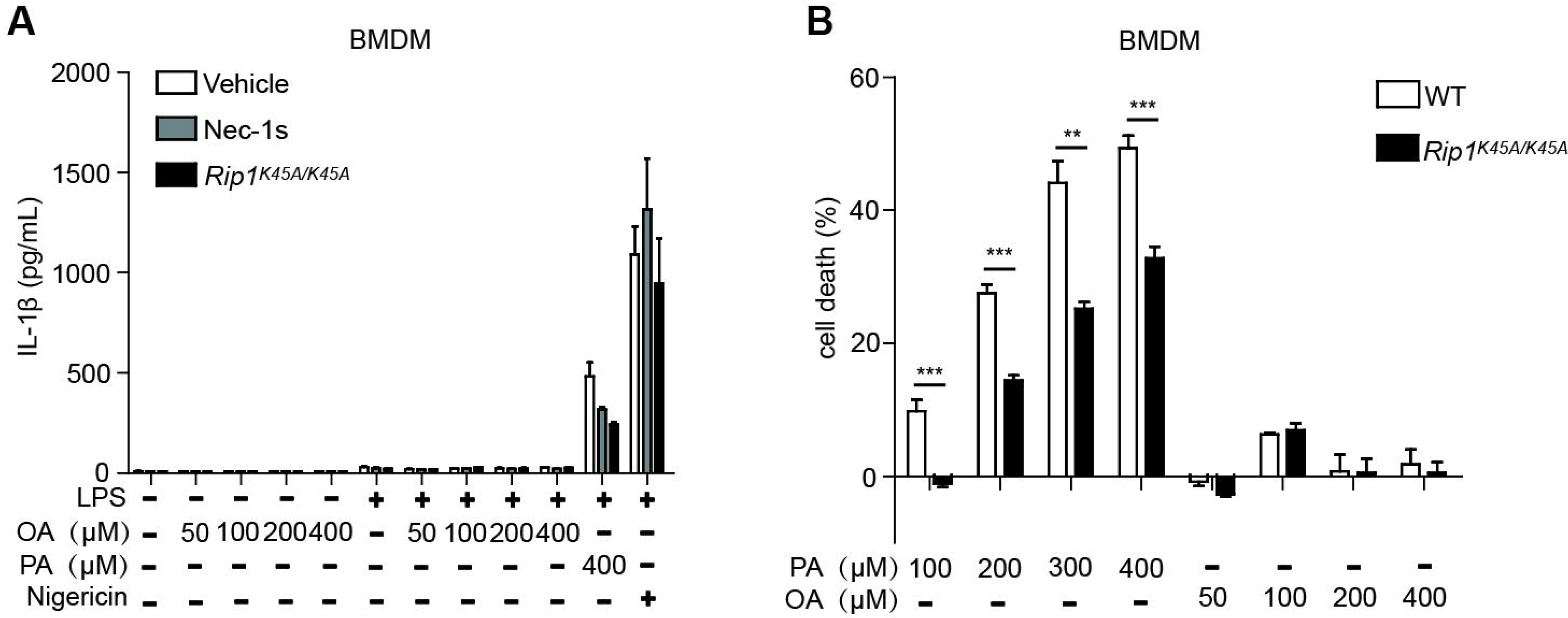
Unsaturated fatty acid oleic acid (OA), unlike saturated fatty acid PA, did not induce obvious cell death and inflammasome activation in BMDMs. BMDMs were treated with different doses of OA with or without LPS priming as indicated for 24 h. Then IL-1β release and cell death rate were detected by ELISA or LDH release assay respectively. Data are expressed as mean ± SEM. ns not significant, *\*p*<0.05, *\*\*p*<0.01, \*\*\**p*<0.001.

**Supplementary Figure 5.**
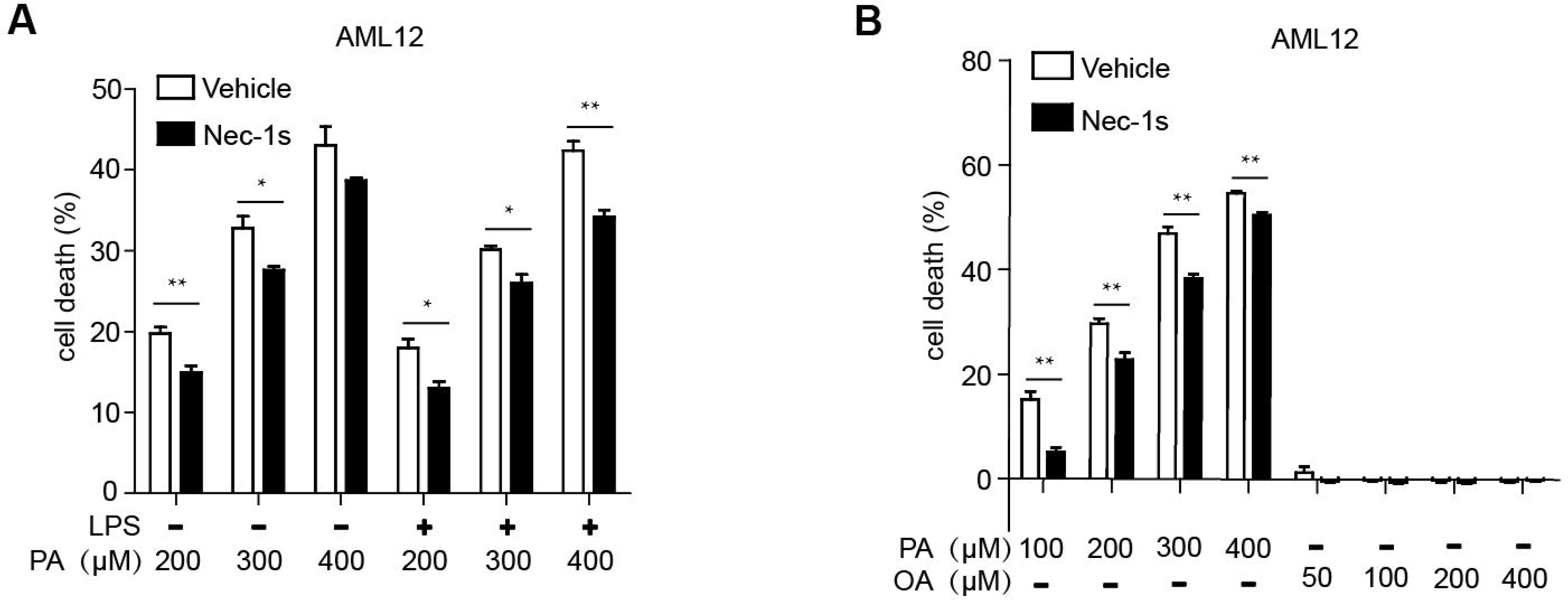
Saturated fatty acid PA, but not unsaturated fatty acid OA, induced cell death in AML12 mouse hepatocytes in a RIP1 kinase-dependent manner. After pretreatment with Nec-1s or vehicle control, AML12 cells were administered by different concentrations of PA, LPS plus PA or OA for 24 h. Then cell death was examined by LDH release assay. Data are expressed as mean ± SEM. ns not significant, *\*p*<0.05, *\*\*p*<0.01, \*\*\**p*<0.001.

**Supplementary Figure 6.**
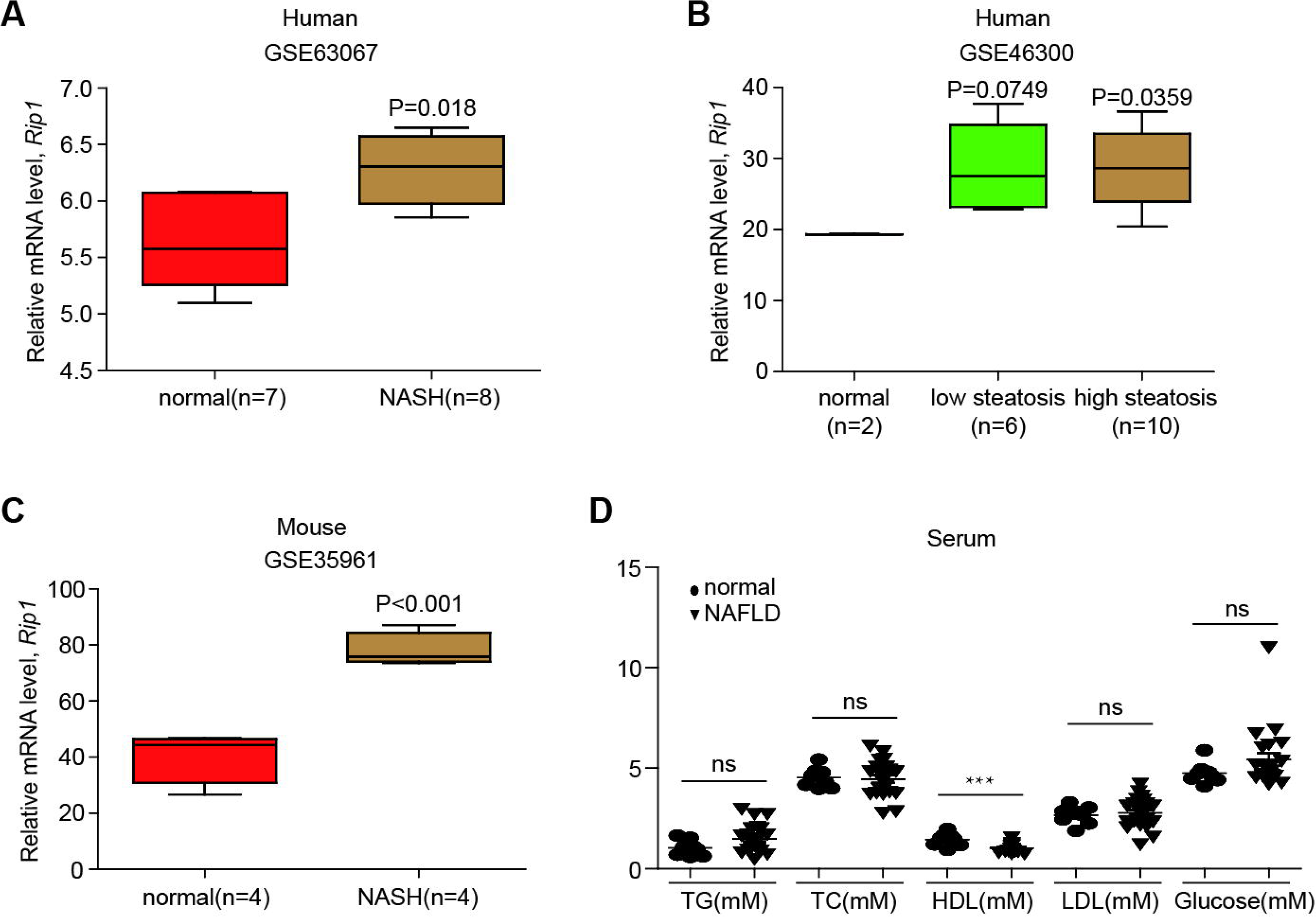
Expression of RIP1 was increased in liver tissues of NASH patients or mice. (A-B) Analysis of mRNA levels of RIP1 using different public datasets to compare RIP1 expression between healthy controls and NASH patients. (C) Analysis of RIP1 mRNA levels in normal or NASH mice using public dataset. (D) The serum levels of TG, TC, HDL, LDL and glucose of the normal persons or NAFLD patients summarized in Table 1. Data are expressed as mean ± SEM. ns not significant, *\*p*<0.05, *\*\*p*<0.01, \*\*\**p*<0.001.

**Supplementary Table 1. Characteristics of the subjects with NAFLD or NASH and healthy controls.**

